# Comparative Genomics of Three Novel Lytic Jumbo Bacteriophages Infecting *Staphylococcus aureus*

**DOI:** 10.1101/2020.12.14.422802

**Authors:** Abby M. Korn, Andrew E. Hillhouse, Lichang Sun, Jason J. Gill

## Abstract

The majority of previously described *Staphylococcus aureus* bacteriophages belong to three major groups: P68-like *Podoviridae*, Twort-like or K-like *Myoviridae,* and a more diverse group of temperate *Siphoviridae*. Here we present three novel *S. aureus* “jumbo” phages: MarsHill, Madawaska, and Machias. These phages were isolated from swine production environments in the United States and represent a novel clade of *S. aureus Myoviridae* that is largely unrelated to other known *S. aureus* phages. The average genome size for these phages is ~269 kb with each genome encoding ~263 predicted protein-coding genes. Phage genome organization and content is most similar to known jumbo phages of *Bacillus*, including AR9 and vB_BpuM-BpSp. All three phages possess genes encoding complete viral and non-viral RNA polymerases, multiple homing endonucleases, and a retron-like reverse transcriptase. Like AR9, all of these phages are presumed to have uracil-substituted DNA which interferes with DNA sequencing. These phages are also able to transduce host plasmids, which is significant as these phages were found circulating in swine production environments and can also infect human *S. aureus* isolates.

**Importance of work:** This study describes the comparative genomics of three novel *S. aureus* jumbo phages: MarsHill, Madawaska, and Machias. These three *S. aureus Myoviridae* represent a new class of *S. aureus* phage that have not been described previously. These phages have presumably hypermodified DNA which inhibits sequencing by several different common platforms. Therefore, not only are these phages an exciting new type of *S. aureus* phage, they also represent potential genomic diversity that has been missed due to the limitations of standard sequencing techniques. The data and methods presented in this study could be useful for an audience far beyond those working in *S. aureus* phage biology. This work is original and has not been submitted for publication in any other journal.

## Introduction

*Staphylococcus aureus* is an opportunistic pathogen of both humans and animals, and is a leading cause of bacteremia, skin, soft tissue and device-related infections in humans (1). The expansion and prevalence of methicillin-resistant *S. aureus* (MRSA) imposes a significant burden to the health care system (2). *S. aureus* infections, particularly MRSA infections, can be difficult and costly to treat, with one study reporting the median cost for treatment of a MRSA surgical site infection as $92,363 (3).

Carriage of *S. aureus* in the general public in the continental US ranges from 26% to 32% (4). An estimated 1.3% of that *S. aureus* being MRSA (5). However, in individuals in the US that are swine farmers, production workers or veterinarians, carriage of multi-drug resistant *S. aureus* (MDRSA) is two to six times greater than individuals in the community, or those who are not exposed to swine (6, 7). As with humans, *S. aureus* is considered to be part of the normal bacterial flora of swine (8). Certain activities on swine farms such as pressure washing and tail docking generate particle sizes capable of depositing primarily in human upper airways but also the primary and secondary bronchi as well as terminal bronchi and alveoli (9). Livestock-associated methicillin-resistant *S. aureus* (LA-MRSA) isolates have been found to have a half-life of five days in settled barn dust with an approximate die-off of 99.9% after 66-72 days (10). Therefore, a better understanding of the ecology of *S. aureus* in these environments and mitigation of MRSA in swine production facilities would be of benefit to farmers and workers for safety reasons, as MRSA isolates are able to persist and possibly spread throughout the environment and workers.

Bacteriophages (phages) are viruses that infect bacteria and are the most abundant organism on Earth with an estimated 10^31^ in the biosphere (11). However, despite their abundance and possible utility there are still many unknowns about the basic biology of most phages and their interactions with their hosts in the environment. Previous investigations of *S. aureus* phages have described three common classes of phages: small, virulent P68-like *Podoviridae* with genomes of ~18-20 kb, various temperate *Siphoviridae* with genomes of ~45 kb, and large, virulent Twort-like *Myoviridae* with genomes of ~130 kb (12, 13). There is a single report of a novel large *S. aureus Myoviridae* that appears to be distinct from the K-like *Myoviridae*; this phage was named S6 as reported by Uchiyama *et al*., in 2014 (14). While S6 was estimated to have a 270 kb genome by pulse-field gel electrophoresis and contain DNA in which thymine was replaced by uracil, no genomic sequence for S6 has been reported. Phages that have genomes over 200 kb are classified as “jumbo” phages (15). Most jumbo phages have been isolated from Gram-negative hosts, with the only known jumbo phages for Gram-positive hosts infecting *Bacillus* spp. Jumbo phage phylogeny and taxonomy are complicated by their often distant relationships to each other and to other known phage types; jumbo phage genomes typically encode large numbers of hypothetical proteins (15).This study describes three novel *S. aureus* jumbo phages isolated from swine environments: MarsHill, Machias and Madawaska.

## Methods

### Culture and maintenance of bacteria and phages

*S. aureus* was routinely cultured on trypticase soy broth (Bacto TSB, Difco) or trypticase soy agar (TSA, TSB + 1.5% w/v Bacto agar, Difco) aerobically at 30 °C. Phages were cultured using the double-layer overlay method (16) with 4 ml of top agar (10 g/L Bacto Tryptone (Difco), 10 g/L NaCl, 0.5% w/v Bacto agar, (Difco)) supplemented with 5 mM each CaCl_2_ and MgSO_4_ over TSA bottom plates. Lawns were inoculated with 0.1 ml of a mid-log *S. aureus* bacterial culture grown to an OD_550_ of ~0.5. Phage stocks were produced by the confluent plate lysate method (17) using the original phage isolation host and harvested with 4-5 ml of lambda diluent (100 mM NaCl, 25 mM Tris-HCl pH 7.4, 8 mM MgSO_4_, 0.01% w/v gelatin). Harvested lysates were centrifuged at 10,000 x g, 10 min, 4 °C, sterilized by passage through a 0.2 μm syringe filter (Millipore, Burlington, MA) and stored in the dark at 4 °C. Phages that could not be cultured to high titers using the double-layer overlay method were propagated in liquid culture by inoculating TSB 1:100 with an overnight culture of the appropriate *S. aureus* host strain and incubating at 30 °C with shaking (180 rpm) until OD_550_ of ~0.5 was reached. The culture was inoculated with phage at a multiplicity of infection (MOI) of 0.01 and incubated at 30 °C with shaking (180 rpm) overnight. The next day the lysate was harvested by centrifugation (10,000 x g, 10 min, 4 °C) and the supernatant was sterilized by passage through a 0.2 μm syringe filter (Millipore).

### Phage isolation

Phages were isolated from environmental swabs collected from 2015 - 2017 in swine production facilities in the United States. Five swabs per site were collected, targeting areas in production facilities with visible residue such as floor slats or water lines. Swabs were collected dry or were wetted in Stuart’s Medium (BD BBL CultureSwab) for transport. Swab heads were aseptically clipped into a 50 ml conical tube (Falcon Corning) and eluted in sterile TSB with shaking for 2 hours at room temperature. Swabs were pooled by collection site. The swab eluates were centrifuged at 8,000 x g, 10 min, at 4 °C, and the supernatants passed through 0.2 μm syringe filters (Millipore). Samples were enriched for phage using a mixed enrichment approach (18, 19)by mixing 12 ml sample supernatant with 8 ml TSB, inoculating with 0.2 ml of a mixed *S. aureus* culture, and incubating overnight at 30 °C with aeration. Enrichment inocula were prepared by mixing equal volumes of overnight *S. aureus* cultures. The sample collected from the O.D. Butler, Jr. Animal Science Complex (College Station, TX) was enriched with a mixture of four *S. aureus* strains: USA300-0114, HT20020371, HT20020354 and CA-513 (**Table 1**). All other samples were enriched using two separate mixed enrichment panels: one panel consisted of a mixture of four human-associated *S. aureus* isolates and the other contained six swine-associated isolates (**Table 1**). Enrichment cultures were centrifuged (8,000 x g, 10 min, 4 °C) and the supernatants sterilized by passage through a 0.2 μm syringe filter (Millipore) and stored at 4 °C. Samples were screened for the presence of phage by spotting 10 μl aliquots onto soft agar lawns inoculated with each individual *S. aureus* strain used for enrichment; samples were scored as positive for phage if they produced clearing zones or visible plaques on a lawn of any *S. aureus* host. Phages were isolated from phage-positive samples by dilution and plating to lawns of *S. aureus* followed by picking of well-isolated plaques. Each phage was subcultured three times to ensure clonality.

**Table 1.** *S. aureus* strains used in this study.

| Strain No. | Strain Name | ST | CC or PFGE | SCCmec | Geographic Origin | Isolation |
| --- | --- | --- | --- | --- | --- | --- |
| NRS384 | USA300-0114 | 8 | USA300 | IV | Mississippi, USA | Human |
| NRS255 | HT20020371 | 80 | 80 | IV | France | Human |
| NRS253 | HT20020354 | 398 | 398 | NONE | France | Human |
| NRS653 | CA-513 | 5 | USA800 | IV | California, USA | Human |
| N/A | PD6 | 9 | N/A | NONE | Illinois, USA | Swine |
| N/A | PD10 | 398 | N/A | YES | Iowa, USA | Swine |
| N/A | PD17 | 398 | N/A | NONE | Texas, USA | Swine |
| N/A | PD18 | 9 | N/A | NONE | North Carolina, USA | Swine |
| N/A | PD19 | 5 | N/A | NONE | North Carolina, USA | Swine |
| N/A | PD32 | 9 | N/A | NONE | Alabama, USA | Swine |

### Phage DNA purification and sequencing

High titer (>10^8^ PFU/mL) lysates of each phage were produced from plate lysates or liquid cultures as described above, and gDNA was extracted from 10-20 ml of phage lysate. Phage genomic DNA was extracted using the Wizard DNA Cleanup Kit (Promega) by a modified protocol as described previously (20, 21). Phage DNA was sheared to fragments of approximately 350 bp using a Diagenode Bioruptor® Pico (Diagenode Diagnostics). One μg of DNA for each phage isolate was used for Illumina Truseq library preparation using the manufacturer’s PCR-free library preparation protocol. Prepared libraries were quantified using KAPA qPCR library quantification kit (Roche) and diluted to 4 nM. Sequencing of phage DNA was performed by Illumina MiSeq V2 Nano 500 cycle chemistry. Library preparation and sequencing were performed at The Institute for Genomics and Society core laboratory at Texas A&M University.

FastQC (22), and SPAdes 3.5.0 (23) were used for read quality control and read assembly, respectively. Analysis of phage contigs by PhageTerm (24) indicated a circularly permuted genome. MarsHill termini were determined by PCR using primers (forward 5’-AGCAGGTATTACAGGCCATTT-3’) (reverse 5’-GAAGACGAAGTTAATGAGGCTAGA-3’) facing off the contig ends to produce a ~1000 bp product. DNA polymerase Phusion U (Thermo Fisher) was used in this end closure PCR and the PCR product was sequenced by Sanger sequencing to close and correct the MarsHill contig. The Madawaska and Machias contigs were closed by reassembling the genomes with SPAdes (23) to produce contigs in which the original contig ends were internal to the genome with >40X coverage; this information was used to correct and close these contigs.

### Restriction digests of phage DNA

To distinguish different phage types approximately 300 ng of extracted gDNA was digested with both DraI (5’ TTTAAA 3’), EcoRI-HF (5’ GAATTC 3’) (New England BioLabs Inc., Ipswich, MA). gDNA samples that showed poor banding patterns or could not be digested by the enzymes listed above were then digested with Taq⍺I (5’ TCGA 3’) and MspI (5’CCGG3’). For DraI, EcoRI-HF and MspI 300 ng of gDNA from each phage was incubated with each enzyme and CutSmart ® Buffer at 37 °C overnight. For gDNA treated with Taq⍺I, 300 ng of gDNA was incubated with Taq⍺I and CutSmart ® Buffer at 65 °C for two hours. After incubation, 4 μL of loading dye was then added and the total 24 μL for each sample was run on a 1% agarose gel at 90 V for 2.5 h.

### Phage genome annotation

Genome annotation was conducted on the Center for Phage Technology Galaxy-Apollo genome annotation platform using the Galaxy pipelines described in (25). Briefly, genes were identified using Glimmer v3 (26) and MetaGeneAnnotator v1.0 (27), and tRNAs were identified using ARAGORN v2.36 (28). Protein function annotations were assigned based on results from BLAST v2.9.0 against the nr and SwissProt databases (29, 30) InterProScan v5.33 (31), HHPred (32), and TMHMM v2.0 (33). Genomic DNA sequence was compared to that of other phages using BLASTn against the nt database, and using ProgressiveMauve v2.4 (34).

### Transduction experiments

The ability of phage MarsHill to transduce host DNA into a recipient cell was determined using methods based on those described in Stanczak-Mrozek et al., 2015 (35). *S. aureus* strains Xen 36 (PerkinElmer; containing a plasmid-borne kanamycin resistance marker) and CA-347 (NRS648; a USA 600 MRSA strain with a chromosomal *mecA* marker) were used as donor *S. aureus* strains, and the swine-associated strain PD17 was used as the recipient. Phage MarsHill was propagated on donor strains in liquid culture as described above. PD17 was grown in 10 ml TSB supplemented with 5 mm MgCl_2_ at 37 °C to ~5×108 CFU/ml. MarsHill lysate propagated on a donor strain was added to the recipient culture at an MOI of 0.1 and incubated for 20 minutes at 30 °C. The culture was centrifuged at 13,000 x g for 2 minutes and the cell pellet was resuspended in 1 ml TSB with 50 mM sodium citrate. This culture was then incubated at 37 °C for 1 h, washed twice in TSB by pelleting and resuspension, and resuspended in 10 ml TSB. This culture was plated in 1 ml aliquots onto TSA plates supplemented with either 150 μg/ml kanamycin (for lysates propagated on Xen 36) or 5 μg/ml oxacillin (for lysates propagated on CA-347) and incubated overnight at 37 °C. Transduction efficiency was calculated as the number of transductants (CFU)/number of input phage particles (PFU), with a detection limit of 2×10^−9^.

### Transmission electron microscopy

Transmission electron micrographs of each phage were obtained by adhering the phage to a carbon film by the Valentine method (36), and staining with 2% uranyl acetate. Phage were viewed in a Jeol 1200EX TEM at 100 kV accelerating voltage.

## Results and Discussion

### Phage isolation and characterization

Phages MarsHill, Madawaska and Machias were isolated from *S. aureus*-enriched environmental samples collected from swine production facilities located in Texas and Minnesota (**Table 2**). MarsHill and Madawaska were isolated against a human clinical USA300 MRSA host, and Machias was isolated on an ST9 MSSA isolate of swine origin. Phage enrichment and culture was routinely conducted at 30 °C, and phages MarsHill, Madawaska and Machias were found to be unable to form plaques or propagate in liquid culture at 37 °C. Temperature sensitivity between 37 °C and 30 °C was also noted for the *S. aureus* jumbo phage S6 (14), and may be a limiting factor for recovery of this type of phage from the environment.

**Table 2.** Summary statistics for three *S. aureus* jumbo myophage genomes.

| Phage Name | *S. aureus* host | Isolation site | Genome Length (bp) | Protein-coding genes | Disrupted genes | Accession no. |
| --- | --- | --- | --- | --- | --- | --- |
| **MarsHill** | USA300-0114 | Texas swine barn | 266,637 | 262 | 9 | MW248466 |
| **Madawaska** | USA300-0114 | Texas swine barn | 265,446 | 264 | 10 | MW349129 |
| **Machias** | PD32 | Minnesota swine barn | 274,478 | 263 | 13 | MW349128 |

Transmission electron microscopy of MarsHill revealed a large phage with typical *Myoviridae* morphology (**Figure 1**). MarsHill was observed to have a mean head size of ~115 nm (± 6 nm) and a tail length of ~233 nm (± 4 nm). For comparison, the K-like *S. aureus* phage phi812 has a reported head diameter of 90 nm and a tail length of 240 nm (37). The measurements of the jumbo phage reported here are comparable to those of phage S6, with a reported head diameter of 118 nm and tail length of 237 nm (14). Although the genome sequence of S6 is not available as of this writing, the large size and temperature-dependent growth characteristics reported for this phage makes it likely to be related to the jumbo phages reported here.

**Figure 1.**
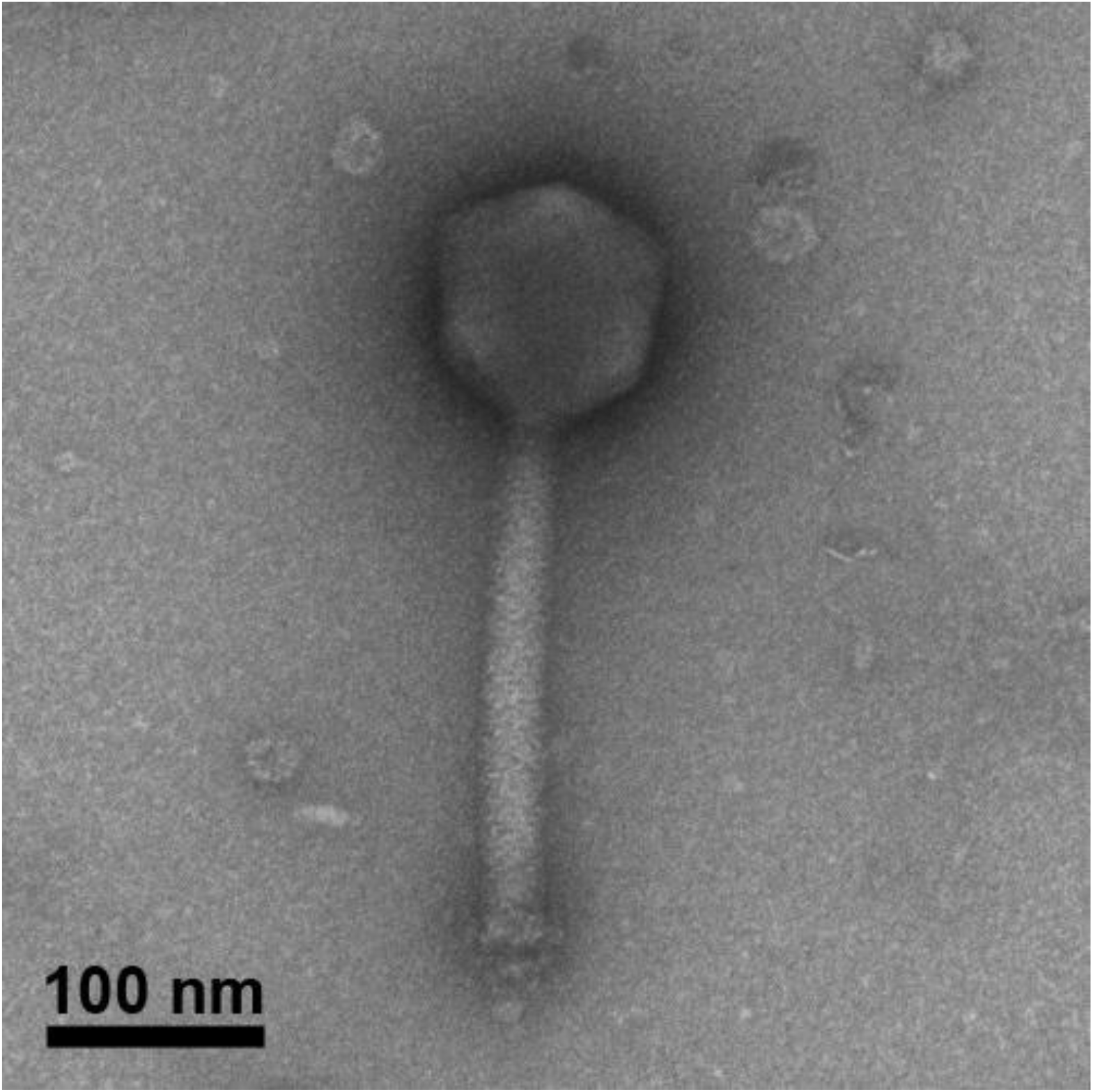
Transmission electron micrograph of the jumbo *S. aureus* myophage MarsHill.

### DNA sequencing and genomic characterization

Genomic DNA extracted from all three phages was not able to be digested by EcoRI-HF (5’ GAATTC 3’) or DraI (5’ TTTAAA 3’), but could be digested with MspI (5’ CCGG 3’) and Taq-αI (TaqI) (5’ TCGA 3’) (**Figure 2**). TaqI was used as it has been shown to cut heavily modified phage DNA such as that of SPO1, which contains hydroxymethyluracil in place of thymine in its DNA (38). These results are consistent with phage DNA modifications to adenine or thymine bases, as MspI, which lacks A or T bases in its recognition sequence, was also able to digest this DNA.

**Figure 2.**
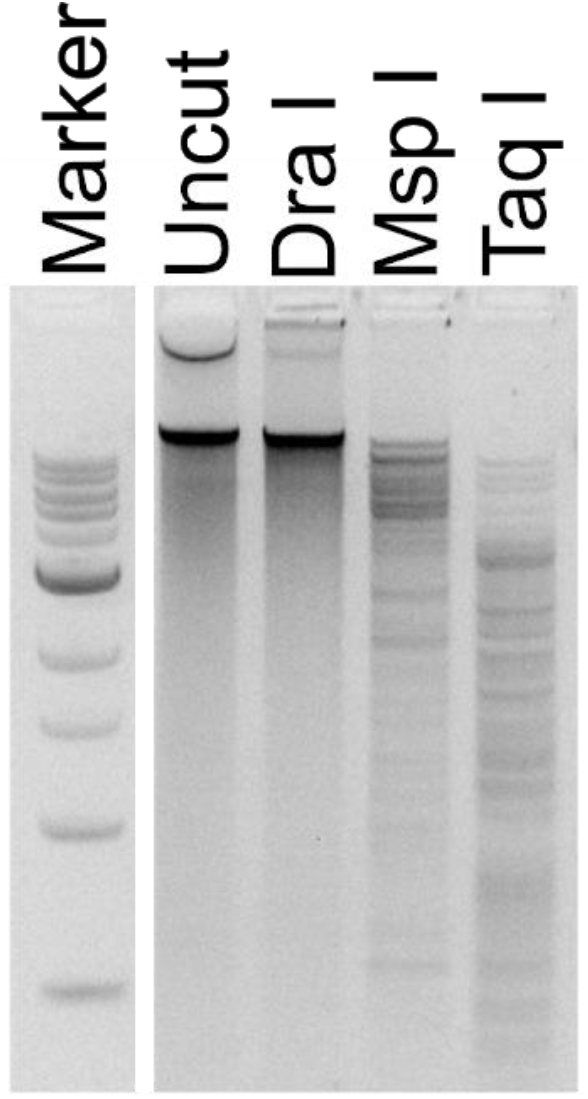
Restriction digests of MarsHill genomic DNA. DNA was not digestible with DraI but was cut with the enzymes MspI and TaqI. Marker is a 1 kb ladder (NEB).

Obtaining the genomic DNA sequence of these phages was challenging. Multiple attempts at standard DNA sequencing approaches using library preparation by Illumina TruSeq Nano or Swift Accel-NGS 1S plus (Swift Biosciences) followed by MiSeq V2 Nano 500 cycle sequencing produced no detectable phage DNA sequence. Pacific BioSciences SMRTbell Express (Pacific Biosciences) failed to produce sequencable DNA libraries. Sequencing by MinION (Oxford Nanopore) produced sequence data that could not be interpreted, even though the DNA sequence of phage lambda could easily be recovered as the positive control for the sequencing run. Illumina library preparation by a PCR-free library protocol, which required significantly higher amounts of input DNA, produced DNA libraries sequencable by Illumina MiSeq. Modification or base substitution of phage DNA has been reported as an impediment to sequencing for other phages, such as the 5-hydroxymethyluracil substitution for thymine found in the *Bacillus* phage CP-51 (39), or uracil substitution for thymine in the *Bacillus* jumbo phage AR9 (40). Phage AR9 was sequenced by Illumina following library amplification using a uracil-compatible DNA polymerase (40) and CP-51 was only sequencable by a combination of PacBio and Sanger sequencing (39). The previously reported *S. aureus* jumbo phage S6 was also determined to contain DNA with thymine completely replaced by uracil (14). The behavior of the DNA of the MarsHill-like phages reported here: resistance to digestion by restriction enzymes with A/T bases in the recognition site, sensitivity to digestion by TaqI, and inability to be amplified by PCR using polymerases other than the uracil-insensitive PhusionU, is consistent with such a thymine-uracil base substitution as reported for phages AR9 and S6 (14, 40).

The substitution of non-standard bases in phage DNA is likely an adaptation that provides broad protection of the phage chromosome from cleavage by host restriction systems (41). The glycosylated 5-hydroxymethylcytosine (glc-HMC) of coliphage T4 protects phage DNA from not only HMC-specific nucleases but also CRISPR-Cas9 (42). Substituting uracil for thymine is also a means of circumventing host defenses; *B. subtilis* phage SPO1 has hydroxymethyluracil in place of thymine, and replacement of this base with thymine renders the phage DNA more susceptible to restriction enzymes (43, 44). It is worth noting that the incompatibility of such hypermodified phage DNA with standard sequencing approaches likely impacts various metaviromic studies of natural phage communities, as this DNA will be rendered “invisible” to such approaches unless procedures are adapted to capture this sequence.

### Genomic analysis of MarsHill-like phages, a clade of phages related to *Bacillus* jumbo phages

Phages MarsHill, Machias and Madawaska were sequenced by Illumina MiSeq to final coverages of 271-, 151- and 212-fold, respectively. These phages are “jumbo” phages with an average genome size of ~269 kb (**Table 2**). The phages’ G+C content of ~25% is lower than the median G+C content of 32.7% found in their *S. aureus* host. Comparison of phage DNA sequences by progressiveMauve (34) (**Figure 3**), found MarsHill and Madawaska were more closely related (92.4% identity), with the Machias DNA sequence more diverged (61.5% identity with MarsHill, and 61.8% identity with Madawaska). In addition to being ~8 kb longer than either MarsHill or Madawaska, Machias contains an inversion in its genome that corresponds to MarsHill genes *171* to *193*. Differences between these genomes are primarily confined to loss or replacement of individual genes, typically in predicted homing endonucleases or hypothetical genes (**Figure 3**). A blastclust comparison of the 789 predicted proteins found across the three *S. aureus* jumbo phages (threshold of >50% identity aligned over >50% protein length) generated 311 protein clusters, with 197 of these conserved in all three phages. Sixty-two of these protein clusters were singletons found in only a single phage, with the majority (46/62, 74%) found only in Machias. The results of the protein cluster analysis are provided in **Supplementary Table S1**.

**Figure 3.**
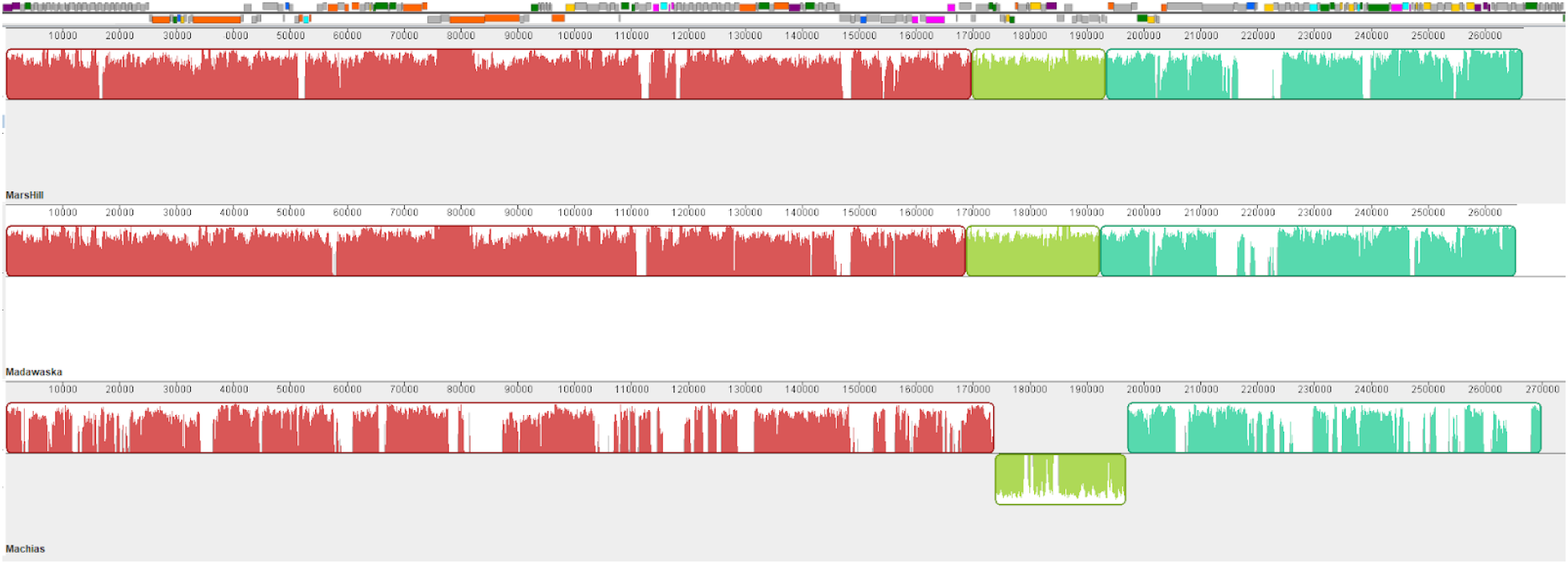
Comparison of three *S. aureus* jumbo phage genomes by progressiveMauve. The annotated genome map for phage MarsHill is added to scale at the top of the figure. The genomes are generally syntenic, with the exception of an inverted region in phage Machias. DNA sequence discontinuities are primarily located at homing endonuclease genes (light blue) or hypothetical genes (grey).

There is little detectable DNA sequence similarity between the MarsHill-like phages and any other organisms in the NCBI database (<5% alignable DNA sequence identity). Comparison of the predicted MarsHill, Madawaska and Machias proteomes to other phages by the CPT Galaxy comparative genomics workflow (25) shows that these phages are most closely related to previously described *Bacillus* jumbo phages AR9 (NC_031039 (32)) PBS1 (NC_043027), and vB_BpuM-BpSp (KT895374.1 (45)) (**Table 3**). These phages are more distantly related to jumbo phages phiR-37 and MS32 infecting Gram-negative hosts *Yersinia enterocolitica* and *Vibrio mediterranei*, respectively. Phage Machias shares a greater number of proteins with these other jumbo phages, and also shares more proteins with the staphylococcal myophage Twort and other Twort-like phages (**Table 3**). Genes shared between the MarsHill-like and Twort-like phages are primarily those encoding homing endonucleases (14 of 21 shared proteins), ribonucleotide reductases (2 proteins) and a predicted amidase. The AR9-like jumbo phages are part of a larger clade of phiKZ-like jumbo phages that can be found infecting Gram-negative and Gram-positive hosts (38).

**Table 3.** Predicted proteins shared between the MarsHill-like phages and other closely related phages, as determined by BLASTp (E<10^−5^). Top row lists the related phage name, NCBI accession and phage host.

|  | AR9<br>NC_031039<br>*B. subtilis* | PBS1<br>NC_043027<br>*B. subtilis* | BpuM-BpSp<br>KT895374<br>*B. pumilus* | phiR-37<br>NC_016163<br>*Y. enterocolitica* | MS32<br>MK308677<br>*V. mediterranei* | Twort<br>NC_007021<br>*S. aureus* |
| --- | --- | --- | --- | --- | --- | --- |
| **Mars Hill** | 77 | 76 | 75 | 60 | 20 | 13 |
| **Madawaska** | 76 | 75 | 75 | 59 | 19 | 12 |
| **Machias** | 81 | 81 | 80 | 62 | 19 | 21 |

The genomes of phages MarsHill, Madawaska and Machias were annotated using the CPT Galaxy-Apollo toolset (25). Due to the overall similarity of the three phages, they will be discussed primarily using phage MarsHill as a reference. The phage genomes were closed and then analyzed by PhageTerm (24) to determine genomic termini, which indicated phage chromosomes with *pac*-like circular permutation. The phage genomes were reopened immediately upstream of the genes encoding predicted non-viral RNA polymerase β and β’ subunits N-terminal domains (**Figure 4**). The MarsHill genome encodes 262 predicted proteins, of which 76 could be assigned functions. The genomes exhibit a general lack of clear modular organization, which is common in jumbo phages (15). Multiple structural components could be identified bioinformatically in the MarsHill genome, based on the presence of conserved domains or their similarity to proteins previously described in phages AR9 (40, 46) and vB_BpuM-BpSp (45, 47), including the major capsid protein, predicted tail fibers, tail sheath, baseplate wedge, prohead scaffold and portal protein (**Table S1**). Two potential tape measure proteins were identified; both contain significant alpha-helical content as well as several predicted transmembrane domains which is indicative of tape measure proteins (48), but one contains a detectable peptidoglycan-degrading domains (PF18013, PF00877), and thus was annotated as a potential tail lysin. All three phages contain dCMP deaminase genes, but other genes associated with modifications to uracil such as dUMP hydroxymethylase or HMdUMP kinase were not identified in the genomes of the *S. aureus* jumbo phages (44). A GroEL-like chaperonin protein was identified in all three *S. aureus* jumbo phage genomes; a similar GroEL-like protein has also been identified in phage AR9 (40). The GroEL-like protein in AR9 has been shown to possess chaperone activity without requiring a co-chaperonin to function when purified and expressed in *E. coli* (49). In coliphage T4, a GroEL homolog (T4 gp31) is essential for folding of the major capsid protein and aids in the assembly of phage particles (50), and this jumbo phage chaperonin likely plays a similar role in folding key viral proteins.

**Figure 4.**
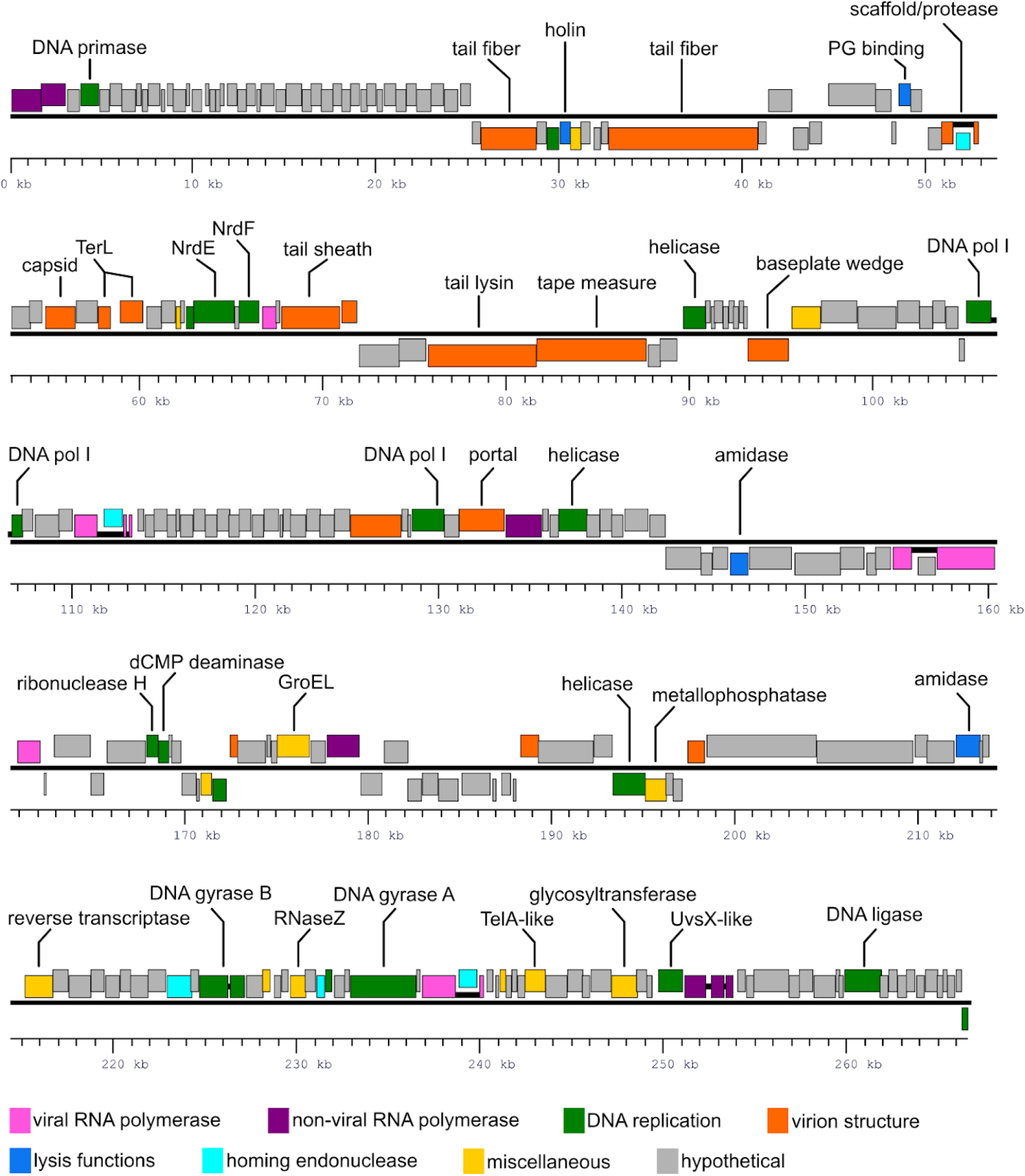
Annotated genome map of the jumbo *S. aureus* phage MarsHill. Gene transcribed from the top DNA strand are drawn above the black line, and genes transcribed from the minus strand are drawn below. Genes are color coded based on their predicted functions as shown in the legend, and selected genes are annotated with their predicted functions. Open reading frames that are predicted to be part of intron-disrupted genes are joined by heavy black lines.

MarsHill gp047 was annotated as the phage holin due to the presence of conserved domains associated with staphylococcal and streptococcal phages (pfam04531, TIGR01598), a conserved protein domain for the holin of lactococcal bacteriophage phi LC3 (51). However, two MarsHill proteins (gp153, gp203) possess characteristics of phage endolysins, thus both proteins have been annotated simply as N-acetylmuramoyl-L-alanine amidases. MarsHill gp203 has similarity to amidases in phages AR9 (40) and PBS1 (NC_043027.1). However, this similarity is to AR9 gp272, not to AR9 gp194, which is noted to have a PlyB-like endolysin conserved domain (cd06523) and is not conserved in the MarsHill-like phages. The second MarsHill amidase, gp153, is more closely related to bacterial amidases and does not share a homolog in AR9. It not clear if one or both of the identified amidases function as the endolysin in the Mars-Hill-like phages; these phages may have “captured” a second amidase gene with redundant endolysin activity.

A large class of phiKZ-like jumbo phages are known to encode their own DNA-directed RNA polymerases (40). These jumbo phage RNA polymerases differ from the bacterial RNA polymerase in that they lack α and ω subunits, with only β and β’ subunits identified (52). These jumbo phages encode two complete RNA polymerase holoenzymes, one of which (the “viral” RNA polymerase) is packaged into the virion and presumably ejected into the host cell upon infection (53), and the other, “non-viral” RNA polymerase expressed during the phage lytic cycle (54). Identification and annotation of the jumbo phage RNA polymerase genes is complicated by their arrangement: each of the viral and non-viral β and β’ RNA polymerase subunits are expressed from multiple loci, encoding subdomains of each of these subunits. The viral RNA polymerase is comprised of β and β’ subunits, with the β subunit expressed from two genes encoding N- and C-terminal domains, and the β’ subunit encoded by three genes encoding N-, mid- and recently-discovered C-terminal domains (54). The non-viral RNA polymerase is comprised of β and β’ subunits, each encoded by two genes corresponding to N- and C-termini, and an additional, fifth protein subunit, which appears to be involved in phage-specific promoter recognition (55). The jumbo phage RNA polymerase subunit genes are also frequently disrupted by one or more introns (40). Genes encoding five viral RNA polymerase subunits (β-N, β-C, β’-N, β’-mid, β’-C) and five non-viral RNA polymerase subunits (β-N, β-C, β’-N, β’-C, specificity determinant) were identified in the MarsHill-like jumbo phages based on the presence of conserved domains and similarity to homologs in related phages.

Comparison of the predicted MarsHill, Madawaska and Machias proteins with their homologs in other phages indicated that multiple genes are interrupted by one or more intron sequences. The presence of intron-disrupted genes is well-known in K-like *S. aureus* phages (56), in the *Bacillus* phages SPO1 and SP82 (57), and the *Bacillus* jumbo phage AR9 (40). As shown in **Table 4**, introns were detected in genes encoding multiple RNA polymerase subunits, the prohead scaffold/protease, DNA polymerase subunits, large terminase, and DNA gyrase B. A similar repertoire of genes is known to be disrupted by introns in AR9 (40). Based on protein sequence alignment to their intact homologs in AR9 and other related organisms, the number of exons in disrupted genes in MarsHill, Madawaska and Machias was predicted (**Table 4**). This approach allowed for the prediction of the mature peptides for all intron-disrupted genes except for the large terminase subunit (MarsHill genes *70* and *71*), which could not be confidently reconstructed based on alignment to any intact protein homolog. Some intron-disrupted genes identified in AR9, such as the non-viral RNA polymerase β’ N-terminal subunit, were found to be intact in MarsHill, while genes that are intact in AR9, such as the DNA polymerase I, were found to be disrupted in the MarsHill-like phages. The predicted number of exons and exon boundary positions was variable between the three *S. aureus* jumbo phages described here, which is consistent with the highly plastic nature of these regions (40).

**Table 4.**
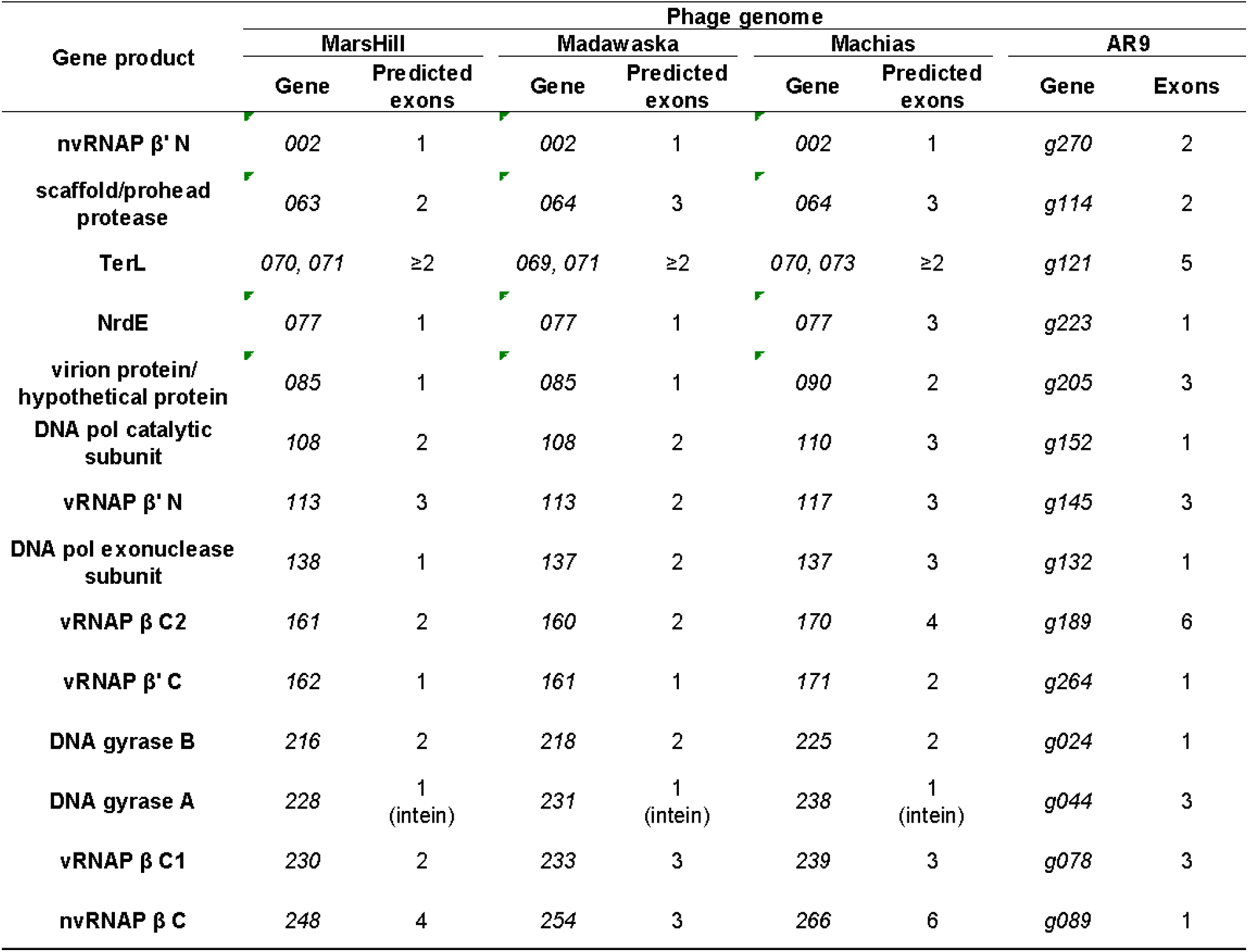
Predicted intron content in the genomes of three *S. aureus* jumbo phages, in comparison with introns identified in the related Bacillus jumbo phage AR9. Only genes disrupted in at least one of the four phages are shown. Intron content in the S. aureus phages was predicted based on protein sequence alignment with intact exons or experimentally determined mature mRNA coding sequences found in AR9 or other related organisms. The DNA gyrase A protein was found to be encoded by a single exon in the MarsHill-like phages but was disrupted by an intein.

Analysis of the protein sequence of the MarsHill DNA gyrase A (gp228) indicated that this gene is disrupted by an intein, based on the presence of intein-associated conserved domains (IPR006142, IPR003587, IPR003586). The intein boundaries in MarsHill gp228 were predicted based on alignment with its intein-less homolog in AR9 (AR9 gp044) and the presence of a conserved cysteine at the N-terminal boundary and the conserved residues His-Asn at the C-terminal boundary (58). Based on this analysis, the intein is predicted to be inserted between residues Y117 and T492 of gp228. The gp228 intein is also predicted to contain a homing endonuclease, based on the presence of a LAGLIDADG endonuclease conserved domain (IPR004860); such domains are a common feature of intein sequences (59). The AR9 homolog of the MarsHill DNA gyrase A appears to lack this intein but is instead disrupted by two intron sequences (40), indicating this gene is a common target for mobile DNA elements.

Previous analysis of AR9 transcription signals identified a AT-rich early promoter that is expressed by the viral RNA polymerase (46). DNA sequence 200 bp upstream of all protein-coding genes was extracted from the MarsHill genome and analyzed for sequence motifs using MEME (60). An AT-rich motif nearly identical to that observed in phage AR9, particularly the consensus motif 5’ TATATTAT 3’, was identified upstream of 29 genes in the MarsHill genome by MEME (**Figure 5**). A second conserved signal corresponding to the phage late promoter was not identified by this approach. Transcriptomic profiling of AR9 has indicated that this signal is somewhat less conserved with a consensus motif of AACA(N_6_)TW (46), and thus is difficult to predict *de novo* without additional data from phage transcripts.

**Figure 5.**
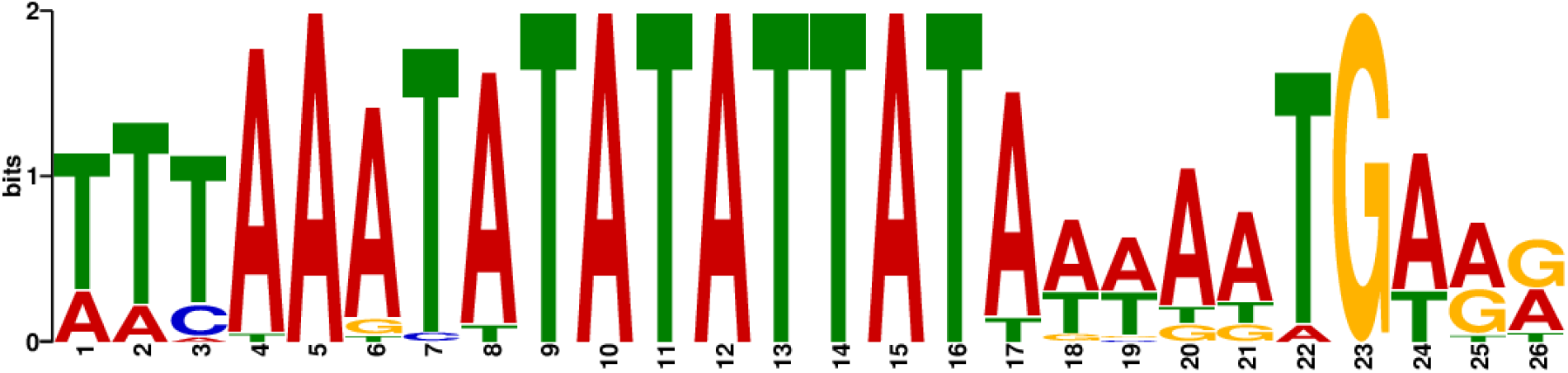
Sequence logo of the predicted MarsHill early promoter sequence recognized by the phage viral RNA polymerase.

All three *S. aureus* jumbo phages encode a predicted RNA-dependent DNA polymerase (reverse transcriptase), exemplified by MarsHill gp206. This 501-residue protein contains a C-terminal retron-like reverse transcriptase domain (cd03487) and a ~200 residue N-terminal domain of unknown function. Reverse transcriptases have been previously identified in phages as part of diversity-generating retroelements (DGR’s) which selectively mutagenize the phage tail fiber to extend phage host range (61, 62). The MarsHill gp206 reverse transcriptase does not appear to be a part of such a system, as the accessory variability determinant (Avd), TR, VR and IMH sequences typically associated with DGR’s were not identified in MarsHill. The MarsHill reverse transcriptase is more closely related to proteins associated with bacterial retroelements, with homologs found in *Bacillus* (TQJ37342, E=10^−26^) and *Marinimicrobium* (ROQ19867, E=10^−21^). The MarsHill reverse transcriptase is quite diverged from other sequences present in NCBI, with the closest bacterial homologs sharing only ~30% sequence identity. Immediately upstream of MarsHill gp206 is a 1,200 bp non-coding region which is hypothesized to contain the retroelement-associated ncRNA (63). Bacterial retroelements have recently been implicated as a type of bacterial anti-phage defense mechanism which leads to abortive infection when an infecting phage inhibits RecBCD, however the MarsHill reverse transcriptase does not appear to be associated with a potential effector protein as described in anti-phage elements (63). If this element serves as an anti-phage system (e.g., to exclude co-infecting phages), it is possible that the gene encoding such an effector has become unlinked in the MarsHill genome. It is also possible that the gp206 reverse transcriptase serves some other function, or is simply present as a selfish genetic element (64).

There are no identified tRNAs in phage MarsHill, with Madawaska having one and Machias two pseudo-tRNA genes as identified by tRNAscan. The presence of tRNAs is positively associated with genome size in phages and is thought to be driven by divergence of phage codon usage from that of the host (65). In these phages, tRNA genes appear to be degenerating, with aberrant pseudo-tRNA genes appearing in two of the phages and these genes being undetectable in MarsHill, indicating a lack of selection pressure to maintain these functions (66).

### Phage MarsHill is able to transduce host DNA

Phages AR9 of *Bacillus* and S6 of *S. aureus* have been previously described as generalized transducing phages, capable of aberrantly packaging host DNA and transferring it to a new cell (40) (14). The ability of MarsHill to transduce chromosomal and plasmid-borne antibiotic resistance marker genes was determined experimentally. MarsHill was able to transduce plasmid-borne kanamycin resistance from *S. aureus* strain Xen 36 to wild-type *S. aureus* isolate PD17 at an efficiency of 7 × 10^−9^ transductants/PFU, based on three replicate experiments. This transduction frequency is comparable to several different *pac*-type *S. aureus Siphoviridae*, which are able to transduce plasmids of varying sizes at a frequency of 10^−5^ to 10^−11^ transductants/ PFU (67). No transductants were recovered using lysates of the virulent *S. aureus* phage K or induced culture supernatants of Xen 36, indicating that any endogenous temperate phages residing in Xen 36 are not produced at high enough levels to account for the observed transduction. Phage K is not expected to be able to perform generalized transduction, as it has been reported to degrade the host chromosome shortly after infection (68), an activity which is associated with lack of transducing capacity in other phages such as the coliphage T4 (69). Transduction experiments conducted with the donor strain CA-347, a USA600 MRSA strain, did not yield any detectable transductants expressing *mecA*-mediated oxacillin resistance in the recipient cultures using either phage or culture supernatants. This observed inability to transduce chromosomal markers is not conclusive as the measured transduction frequency for plasmid DNA was near the detection limit of the assay (2×10^−9^ transductants/PFU), and transduction of the *mecA* locus by temperate *Siphoviridae* phages has been reported ranging from 1 × 10^−9^ to 2 × 10^−10^ transductants/PFU (70). Phage S6 was reported to transduce plasmid pCU1 at an efficiency of 10^−7^ transductants per cell and a plasmid containing the *mecA* gene at an efficiency of 5.2 × 10^−11^ per cell into the restriction-deficient *S. aureus* strain RN4220. S6-mediated transduction of the pCU1 plasmid was also tested for several other staphylococci including *S. epidermidis*, *S. pseudintermedius*, *S. sciuri* and *S. felis*, with reported efficiencies of 10^−7^ to 10^−10^ per cell (14). The transduction efficiency of MarsHill may be expected to be lower than that observed for S6, as the recipient strain used to measure MarsHill transduction is a wild-type *S. aureus* strain recently isolated from the environment which presumably has a functional restriction system, unlike the restrictionless *S. aureus* strain RN4220.

## Conclusion

Three novel *S. aureus Myoviridae* phages isolated from swine environments across the US presented unique challenges in both culturing and sequencing. Multiple commonly used approaches failed to produce usable sequence for these phages, however using a PCR-free library preparation method produced libraries sequencable by Illumina. This study raises questions about the diversity of phages with modified DNA that are missed by metagenomic studies using traditional library preparations to identity phage sequences. Complete genomes were obtained for phages MarsHill, Madawaska and Machias, with all phages having genomes well over 200 kb, classifying them as members of a new jumbo phage lineage in *S. aureus* (15). The *S. aureus* phages in this study were isolated from environmental samples at 30 °C instead of the traditional 37 °C, in hopes of more closely mimicking ambient barn temperatures as well as human nasopharynx temperatures which average around 34 °C (71). Without this adjustment these phages would not have been isolated as they do not form observable plaques or propagate efficiently in liquid culture at 37 °C. These three phages are only distantly related to other known phages, with the most similar being *Bacillus* phage vB_BpuM-BpSp (45) and *Bacillus* phage AR9 (40). All three phages carry identifiable viral and non-viral RNA polymerase subunits, many of which are intron-disrupted. Intron and intein disruption of multiple phage genes was identified, including the viral and non-viral RNA polymerases, DNA polymerase, and DNA gyrase. Mature peptides were predicted for all intron-disrupted genes by alignment to homologs in AR9 and other organisms, except for the phages’ TerL. Unlike phage AR9 all three phages contain an RNA-directed DNA polymerase, the function of which is unclear. MarsHill was able to transduce plasmid DNA at an efficiency of 7 × 10^−9^ transductants/PFU which indicates that these phages, in addition to temperate *S. aureus Siphoviridae,* are also possible vectors for horizontal gene exchange in *S. aureus*. This is particularly relevant as these phages were found to be circulating in swine production environments and are able to infect both swine and human *S. aureus* isolates.

## Acknowledgements

This work was supported by the National Pork Board and Texas AgriLife Research. We would like to thank Dr. H. Morgan Scott and Dr. Andrew Hillhouse for their generosity and time in assisting in the sequencing of these phages. We would also like to thank Dr. Peter Davies for providing swine *S. aureus* isolates and for his help in locating phage sampling sites. We acknowledge the support of the Network on Antimicrobial Resistance in *Staphylococcus aureus* (NARSA) for provision of bacterial isolates.

## Tables and Figures

**Supplementary Table S1.** Related proteins in three *S. aureus* jumbo phages as determined by blastclust with a cutoff of 50% identity over 50% of the protein length. Proteins on the same line were determined to be in the same cluster.

**Provided in attached Excel file.**

